# Genetically Encoded FRET Biosensor for Live-cell Visualization of Lamin A Phosphorylation at Serine 22

**DOI:** 10.1101/2024.05.27.596010

**Authors:** Jian Liu, Qianqian Li, Jinfeng Wang, Juhui Qiu, Jing Zhou, Qin Peng

## Abstract

Extensive phosphorylation at Serine 22 (pSer22) on lamin A is the hallmark of cell mitosis, which contributes to the breakdown of nuclear envelope. In the interphase, the pSer22 lamin A exists in low abundance and involves in mechanotransduction, virus infection and gene expression. Numerous evidence emerges to support lamin A regulation on cell function and fate by phosphorylation. However, live-cell imaging tools for visualizing the dynamics of pSer22 lamin A is yet to be established. Herein, we developed a novel lamin A phosphorylation sensor (LAPS) based on fluorescence resonance energy transfer (FRET) with high sensitivity and specificity. We observed the dynamic lamin A phosphorylation during cell cycle progression in single living cells: the increase of pSer22 modification when cells enter the mitosis and recovered upon the mitosis exit. Our biosensor also showed the gradual reduction of pSer22 modification during cell adhesion and in response to hypotonic environment. By applying LAPS, we captured the propagation of pSer22 modification from inside to outside of the inner nuclear membrane, which further led to the breakdown of nuclear envelope. Meanwhile, we found the synchronous phosphorylation of lamin A and H3S10 at mitosis entry. Inhibition of Aurora B, the responsible kinase for H3S10ph, could shorten the mitotic period without obvious effect on the pSer22 modification level of lamin A. Thus, LAPS allows the spatiotemporal visualization of lamin A phosphorylation at Serine 22 site, which will be useful for elucidating the molecular mechanisms underlying cell mitosis and mechanoresponsive processes.

## Introduction

Lamin A is the main component of lamina that support the integration of nuclear envelope (NE) and highly associated with heterochromatin tethering at nuclear preiphery^1-3^. The protein level of lamin A on nuclear membrane determines different cell fate during differentiation^4-7^. The disassembly and degradation of lamin A is dependent on the phosphorylation events^8^, catalyzed by numerous kinases, like CDK1, PKC and CDK5^9-11^. During mitosis, Ser22 of lamin A is extensively phosphorylated by CDK1 and PKC, inducing rapid breakdown of nuclear envelope^12,13^. In the interphase, lamin A phosphorylation at Serine 22 (pSer22) is maintained at a low level^14^, but dynamically regulated in response to matrix stiffness^15, 16^ and virus infection^17^. Recently, pSer22-lamin A was identified to interact with active enhancer and promote gene expression, which contributes to the upregulation of progeria related risky genes^18^. Moreover, lamin A gets phosphorylated at the DNA double-strand break site by kinase ATR, which eventually induce the nuclear envelop (NE) rupture^19^. Although, pSer22 modification on lamin A shows versatile functionality, it lacks live visualization methods to spatiotemporally track the dynamic of pSer22-lamin A for better mechanistic understanding.

Recently, a lamin A nanobody based FRET biosensor was designed to indicate mechanical forces on nuclear lamin A^20^. The FRET biosensors consisted of two lamin A nanobody inserted with an existing FRET module, which could measure the intermolecular force between adjacent lamin A. The biosensor successfully visualized the variation of mechanical strain of lamin filaments after regulation of nuclear volume, functional LINC complex and actomyosin contractility. However, it cannot visualize whether lamin A is depolymerized or not mediating by pSer22 in respond to mechanical force.

Herein, we designed a novel **l**amin **A p**hosphorylation **s**ensor (LAPS) based on fluorescence resonance energy transfer (FRET) to real-time visualize pSer22 modification on lamin A. By screening the lamin A targeting strategy and linker length, we obtained a biosensor that can indicate the pSer22 level on lamin A with high specificity and sensitivity. Using LAPS, we successfully observed the boost of pSer22 modification before the NE breakdown at the entrance of mitosis. Meanwhile, we observed gradual reduction of pSer22 modification during the cell adhesion and in response to hypotonic stimulation, which may correspond to the enhanced lamina stress. With high spatiotemporal resolution of LAPS, we found that pSer22 modification on lamin A that is located on NE occurred gradually from the inside to outside of the inner nuclear membrane when cells enter the mitosis. Concurrently with pSer22 modification on lamin A is the phosphorylation of Serine 10 on histone 3 (H3S10p) and chromatin condensation. When Aurora B, the kinase of H3S10p, was inhibited, the pSer22 modification on lamin A still happened although the mitosis period was shortened, which indicated that phosphorylation on lamin A was relatively independent from Aurora B.

## Materials and Methods

### Plasmid construction

Lamin-A S22 FRET biosensor consists of 5 parts: FHA2 domain, YPet, linker (EV linker or 34-mer), ECFP, and full-length Lamin A (or lamin A nanobody) (PMID: 37391402). A flexible linker was used to separate FRET pair YPet/ECFP. We amplified the complete encoding sequence and cloned onto a pSin vector backbone by Gibson assembly (NEB, E2621S). To further examine the specificity of the biosensor, anergic mutated biosensor R605A and lamin A mutant biosensors S22A/S392A were constructed. A point mismatch on primer sequence was introduced into the coding sequence of FHA2 or lamin A by PCR, then clone to pSin backbone by Gibson Assembly. The final constructs were verified by Sanger sequencing at Genewiz. All the primers used in this study were listed in **Table S1**.

### Cell culture and transfection

HeLa cells were used for biosensor evaluation and analysis in this study. The cells were cultured in DMEM medium that contained 10% fetal bovine serum (FBS) from Sigma-Aldrich and 1% penicillin/streptomycin (Gibco). After that, the cells were grown in a humidified incubator with 5% CO2 at 37 °C. For transfection of pSin-FHA2-Ypet-linker-ECFP-Lamin-A and mutant, the cells were seeded in 24-well plate for 24 hrs, and then Hela cells were transfected with 200 ng plasmid by Lipofectamine™ 3000 Transfection Reagent (Invitrogen). All cells were cultured in the incubator at 37°C with a 5% CO2 in humid air atmosphere.

### Drug treatments

The hypotonic condition was generated by using DMEM culture medium containing 50% water. To get G1/S boundary cells, 2mM thymidine (Sigma, T1895) was added for 18 h in Hela cells. After PBS twice wash, cells were then released into regular medium for 8 h, and then 2mM thymidine treated for another 18h. To arrest cells at G2/M transition, RO-3306 (Beyotime, SC6673-5mg) was added to a final concentration of 9 µm for 18h. To inhibition Aurora B activity, Hesperadin (MCE, HY-12054) was added to a final concentration of 200 nM to the cells. Hesperadin was added 0.5 hr before the imaging with RO-3306 in the medium. When FRET imaging started, cells were released from RO-3306 but with Hesparadin treatment during imaging process.

### Western blot

LAPS WT and mutants were transfected into HeLa cells in a 12-well plate, respectively, using Lipofectamine 3000 (Invitrogen, L3000015) according to the manufacturer’s protocol. After 48 hrs expression, a sharp fluorescent signal can be visualized under confocal microscopy, and at that moment, plain HeLa and transfected HeLa cells were harvested respectively and lysed in RIPA buffer (Thermo, 89900). Cell lysis was centrifugated at 15000 g for 15 min at 4 °C and the supernatant was collected for Western blot analysis. Each sample was stained with three different primary antibodies: lamin A, phosphorylated Lamin-A (pLamin-A), and GFP, sequentially with completed stripping for each staining. The blots were stripped off by Stripping buffer (Thermo, 21059) at 37 °C for one hour before the next round of Ab labeling.

### Immunofluorescence staining

Cells were seeded on the glass-bottom dishes for 12-24h, and then rinsed with PBS for 3 times, then fixed in 4% (w/v) paraformaldehyde for 30 min and then rinsed with PBS for 3 times, permeabilized in 0.25% (v/v) Triton X-100 for 10 min and rinsed with PBS for 3 times. Subsequently, samples were blocked with 5% BSA for 2 h at room temperature. Anti-H3pS10 antibody (1:500, CST, 9706S) and Anti-plamin A antibody (1:500, CST, 13448) were incubated overnight at 4°C and rinsed with PBST (PBS with 0.1% Tween20). Next, the samples were incubated with secondary antibody at room temperature for 2 h, e.g., goat anti-mouse Alexa Fluor® 488 (1:1000, Abcam, ab150077) and Alexa Fluor® 647 (1:1000, CST, 4410S). Finally, samples were rinsed with PBST and incubated with Hoechst 33342 (1:2000, CST, 4082S) for 10 min at room temperature for nuclei staining. After PBST wash, the samples were mounted with VECTASHIELD Antifade Mounting Medium (Vectorlabs, H-1000-10) and took image by Dragonfly Spinning Disk Confocal Microscopy System (Dragonfly CR-DFLY-202-40, U.S.A.).

### FRET imaging and analysis

After 48 hrs of transfection, cells expressing WT LAPS or mutant LAPS biosensors were seeded on the glass-bottom dishes (Biosharp, BS-15-GJM). Before imaging, the cell culture was switched to DMEM medium without phenol red. FRET imaging was conducted on a Dragonfly Confocal Microscopy System equipped with a seven-laser, EMCCD camera, and live-cell workstation. The ECFP and FRET channels were excited by a 445 nm laser. The 478 nm filter and 571 nm filter were used to collect the emission of ECFP and FRET, respectively.

The ECFP and FRET images were processed and quantified by the image analysis software package Fluocell (http://github.com/lu6007/fluocell), and the ECFP/FRET ratio were calculated and analyzed by GraphPad Prism software.

### CRISPR-mediated LMNA gene knockout

To knockout LMNA genes, we first inserted LMNA gRNA into pSpCas9(BB)-2A-GFP (PX458) (addgene#48138). Here we designed primers for two gRNAs (gRNA1: 5’ GCGAGCTCCATGACCTGCGG 3’, gRNA2: 5’ TCTCAGTGAGAAGCGCACAT 3’) and the vectors were assembled by Golden Gate assay. Plasmids were verified by Sanger sequencing. Then, 0.4 million C2C12 cells were seeded into 6-well plate the day before transfection. 1.25 μg LMNA-PX458-gRNA1 and 1.25 ug LMNA-PX458-gRNA2 were co-transfected into C2C12 with Lipofectamine 3000 Reagent (Invitrogen, cat L3000015). 48 hrs post transfection, cells were subjected to FACS (CytoFLEX SRT, Beckman) to isolate GFP positive monoclonal cell into 96-well plates. The knockout efficiency was further verified by genotyping PCR with LMNA-specific primers (LMNA-F: 5’ TCGAGGCTCTTCTCAACTCC 3’, LMNA-R: 5’ AGGTGAGCAGGCAAATGG 3’) as well as Sanger sequencing.

### Statistical analysis

All experiments were conducted in triplicate and repeat at least twice unless otherwise stated. All the data were presented as mean ± standard error (SEM). The significance of differences was analyzed with one-way ANOVA followed by Tukey’s honest significant difference test using GraphPad Prism software (version 8.0, GraphPad Software Inc.). Significant differences were determined by *p*-values (\**p*<0.05, \*\**p*<0.01, \*\*\**p*<0.001).

## Results

### Lamin A phosphorylation sensor (LAPS) design and characterization

To obtain highly sensitive and specific FRET biosensor for pSer22-LaminA detection, we linked the lamin A nanobody and FHA2 domain with two fluorescent proteins ECFP and YPet in the middle. A flexible linker (EV linker) with 120 aa was inserted between ECFP and YPet according to our previous work^21^. This biosensor was named as pLamin A biosensor 1.0 (**Fig S1a**). Different levels of pSer22 on lamin A in interphase cells and mitotic cells were observed by pLamin A biosensor 1.0 with proximately 10% dynamic changes (**Fig S1b-c**), indicating the low sensitivity of pLamin A biosensor 1.0. Considering that the nanobody targeting site is the end of the coiled-coil domain of lamin A, so it is far away from pSer22 site^20^. To increase the specificity to pSer22, we switched the substrate into full-length lamin A based on pLamin A biosensor 1.0, which called pLamin A biosensor 2.0 (**Fig S2a**). Our results showed that the NE location of the biosensor was dramatically improved (**Fig S2b**). The wild-type (WT) biosensor successfully tracked the dynamic changes of lamin A dephosphorylation when cell exited mitosis, while R605A mutation in FHA2 impaired the mitotic response (**Fig S2b-h**). However, the fold change of FRET ratio between mitosis and interphase was still unsatisfied (**Fig S2**). Considering that the FHA2 domain had low affinity with phosphorylated serine^22^, we shortened the EV linker from 120 aa to 34 aa to enhance the possibility of FHA2 association with phosphorylated serine and named it as **l**amin **A p**hosphorylation **s**ensor (LAPS) (**Fig 1a**).

**Fig 1.**
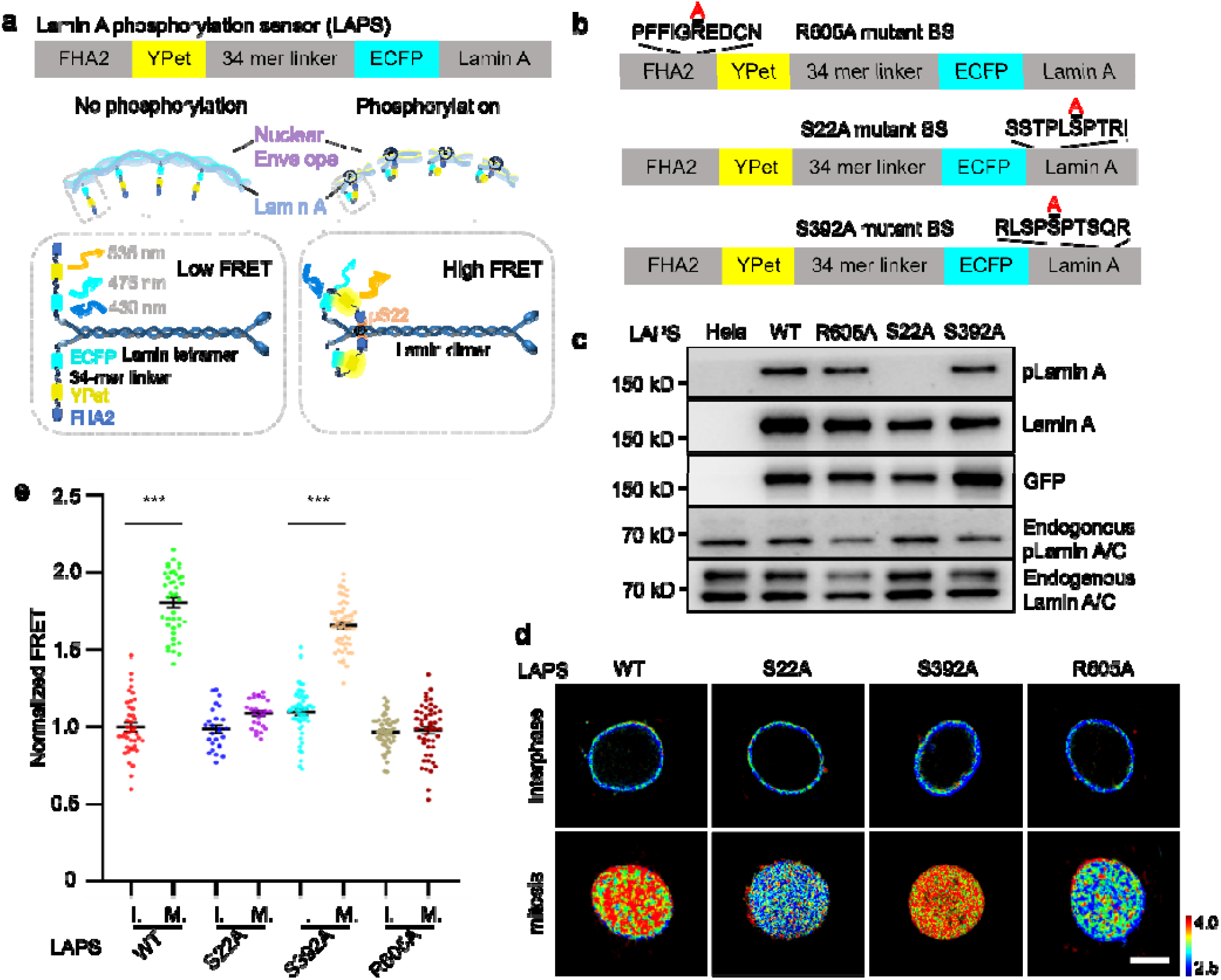
Development and characterization of LAPS. **a**. Schematic diagram of LAPS. At the resting state when lamin A is not phosphorylated, the excitation of ECFP could not trigger a FRET event. When lamin A gets phosphorylated, the FHA2 domain can sense the phosphorylation site, which leads to a tremendous conformational change and proximity of the FRET pair protein. An increase in the FRET ratio could be then observed. Thus, high FRET is corresponding to the phosphorylation state of Lamin A, which mediates depolymerization of lamin A protein. **b**. The construct of WT LAPS and three mutants of LAPS. R605A mutation occurs on the FHA2 domain, while S22A and S392A are on lamin A. **c**. The expression and phosphorylation of LAPS in HeLa cells. All the groups (left to right) represent plain HeLa cells, HeLa cells expressing WT LAPS, R605 LAPS, S22A LAPS, and S392A LAPS. Bands with larger size (∼175 kD) refer to constitutive expression of biosensors. **d**. Representative images of basal level FRET ration from HeLa cells expressing WT, S22A, S392A and R605A LAPS in interphase and mitosis. Scale bar: 10 μm. **e**. Quantitative analysis of the FRET ratio from (d) (n>20). I: interphase, M: mitosis (***P<0.001).

To comprehensively verify the specificity and sensitivity of LAPS, we designed distinct mutations, one was Ser22A in the N terminus of lamin A, the other was Ser392A in the C terminus of lamin A, and another was Arg605 mutated into Ala in the FHA2 domain to get R605A mutation (**Fig 1b**). Upon transient transfection in HeLa cells, we detected lamin A and its phosphorylation level on both biosensors (170 kD) and endogenous lamin A (70 kD) by western blot, and 170 kDa band disappeared in S22A group but not in S392A group, indicating LAPS behaves as endogenous lamin A as we designed (**Fig 1c**). The live cell imaging results showed the correct location of all the biosensors around the nuclear envelope, which indicates the correct expression and function of wildtype and mutated biosensors in mammalian cells (**Fig 1d**). There were similar basal level FRET ratios among WT and mutant biosensors in interphase cells, while much higher FRET ratio in the mitotic cells expressing WT and S392A biosensors due to high pSer22 level during mitosis without changes in S22A and R605A mutant groups (**Fig 1d-e**), implying LAPS detected the pSer22 specifically. In addition, the dynamic range of LAPS was around 80% changes in terms of FRET ratio (**Fig 1e**), which was increased drastically compared to pLamin A biosensor 1.0 and 2.0. Therefore, all the results demonstrated the high specificity and sensitivity of LAPS.

### LAPS detects pSer22 dynamics in cell adhesion and hypotonic condition

It has been shown that the phosphorylation of lamin A changes under mechanical force variation^23^. To visualize the dynamics of pSer22 on lamin A, we transiently transfected Hela cells with WT LAPS and three mutants of LAPS (**Fig 2a-c**). Under the hypotonic conditions, we observed the gradual reduction of FRET Ratio in WT group, which was much lower than S22A and R605A group, indicating that pSer22 on lamin A decreased during the increase of cell tension. Meanwhile, the S392A group changed similarly to the WT group, further implying that our biosensors mainly detected the phosphorylation on Ser22 as designed. The other process monitored by LAPS was cell adhesion. Literatures showed that Lamin A phosphorylation decreases progressively during cell adhesion^15^, which was also observed with our WT LAPS, whereas the R605A mutant showed relatively low response (**Fig 2d-f**). All the results demonstrated that NE adapted to exocellular mechanical change by regulating the pSer22 modification of lamin A and further affecting the balance of lamina polymerization and depolymerization.

**Fig 2.**
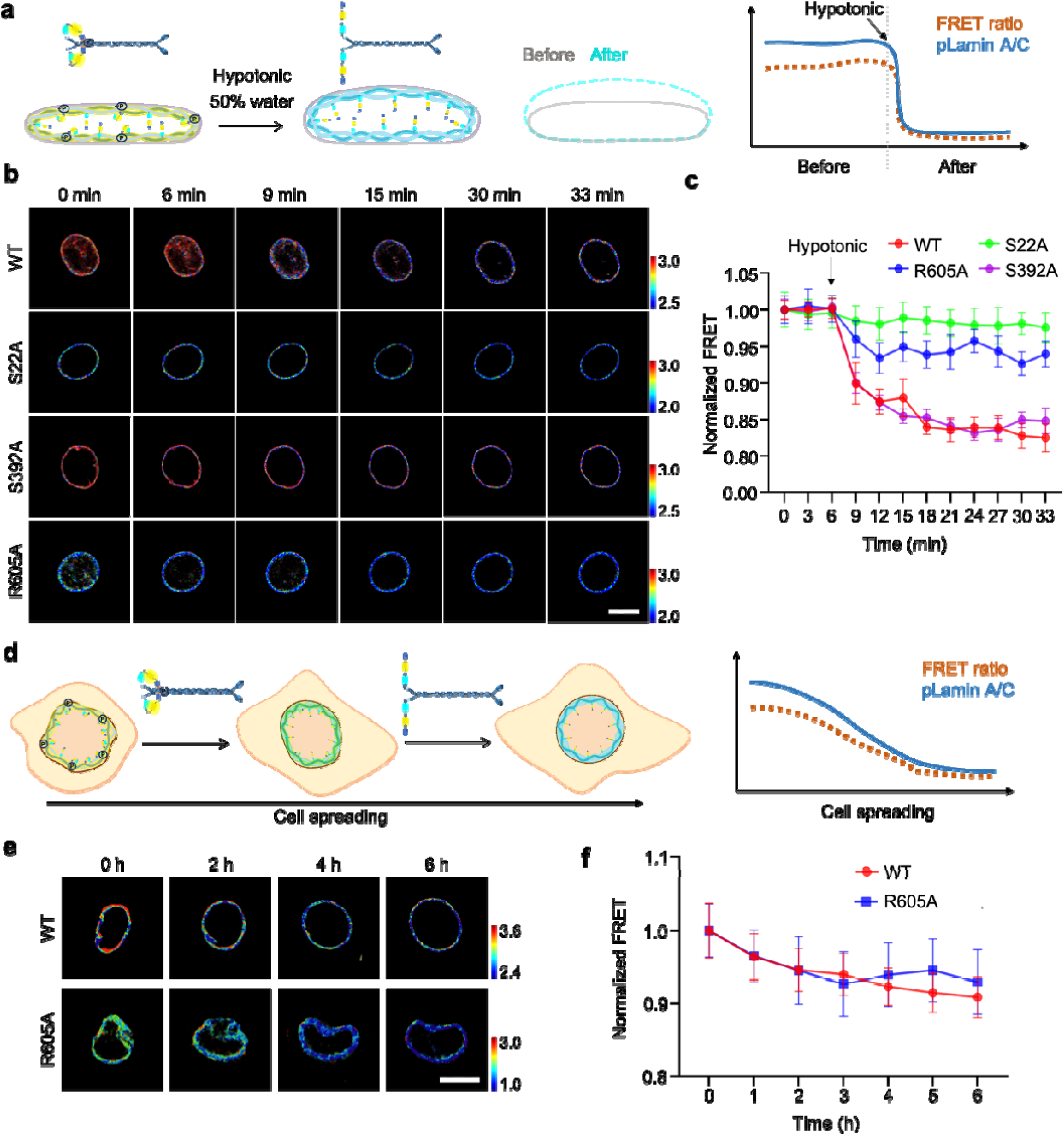
The effect of nuclear deformation on lamin A phosphorylation. **a**. The diagram shows the change in Lamin A phosphorylation at low osmolality. The hypotonic condition was treated by using DMEM culture medium containing 50% water, creating an immediate low osmolality environment in which the nuclei swell and increase the pressure, mediating higher levels of dephosphorylation of lamin A, opening of the FRET biosensor structure, and decreasing FRET ratio. **b**. Time-lapse FRET/CFP ratio images of the WT LAPS and the other three mutants in Hela cells treated with H_2_O in about half-an hour. The color bar indicates high (red) and low (blue) FRET ratio. Scale bar = 10 μm. **c**. Time courses of the normalized FRET/CFP ratio of the WT and mutants (n=20) in Hela cells treated with H_2_O. **d**. The schematic diagram shows the dynamic process of Lamin A phosphorylation during cell spreading. The detached cells are with higher levels of lamin A dephosphorylation, opening of the FRET biosensor structure and decreasing FRET ratio, and vice versa. **e**. Time-lapse FRET/CFP ratio images of the WT LAPS and R605A mutant during Hela cells adhesion onto the matrix. The color bar indicates high (red) and low (blue) levels of pLamin A, scale bar = 10 μm. **f**. Time courses of the normalized FRET/CFP ratio from (e) (n=17).

### LAPS detects pSer22 dynamics in cell cycle

LAPS detected the significant pSer22 modification in mitotic cells compared to interphase cells (**Fig 1d-e**), so we wonder the entire dynamics of pSer22 on lamin A during cell cycle. We firstly synchronized the HeLa cells transiently expressing WT biosensors by CDK1 inhibitor RO-3306 to late G2 phase^24^ and then released the cells to go through the whole mitosis (**Fig 3a**). We found that the NE broke down immediately after removal of Ro--3306. The FRET ratio extensively raised as the NE breakdown occurred (**Fig 3b and Movie 1**). In the telophase, the FRET ratio on the chromosome was reduced, which was consistent with the previous reports that pSer22 modified lamin A was dephosphorylated by the phosphatase PP1 which located on the chromosome surface in the telophase (**Fig 3b**)^25^. The time-course images exhibited the high spatiotemporal resolution of LAPS. To further refine the observation of the changes in pSer22 modification on lamin A during the whole cell cycle, we did more careful quantification of the FRET ratio from interphase released from double thymidine to mitosis or from mitosis to interphase. As the cell progressed from interphase into mitosis, the NE gradually broke down and the FRET signal increased (**Fig 3c-d**). When the mitotic cells entered interphase, the evenly distributed fluorescent signal was gradually concentrated in two regions within the cell representing the reassembled nuclei. And the FRET signal was declined along cell division (**Fig 3e-f**). The decrease in FRET ratio reflects the reduction of pSer22 modification on lamin A, which occurs when lamin A concentrated around the chromosome in nascent nucleus to prepare for reconstruction of nuclear membrane. On the other hand, cells expressing the mutant biosensor R605A lacked the FRET signaling response during mitosis.

**Fig 3.**
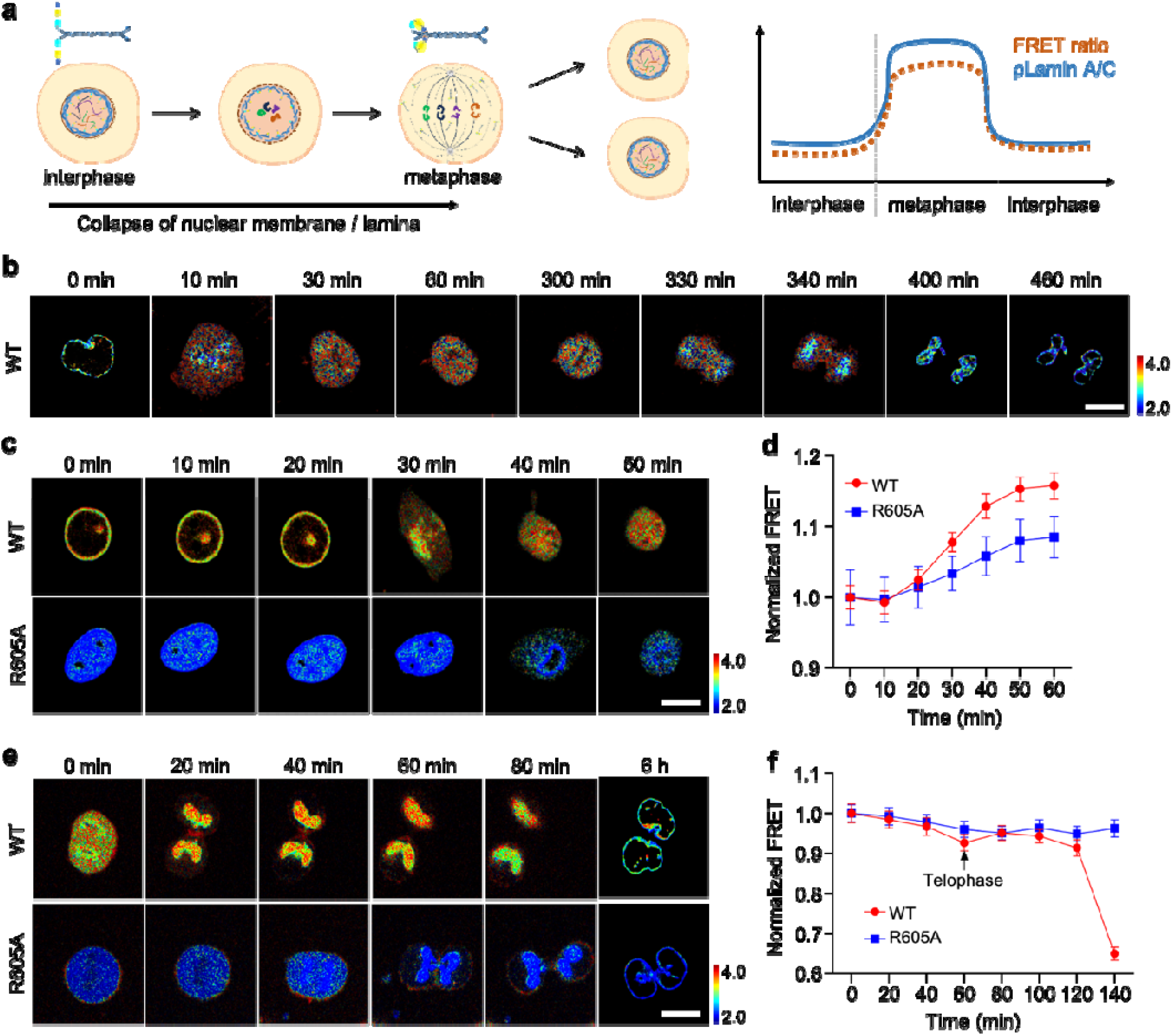
The dynamics of pSer22 on lamin A were monitored by LAPS during cell cycle. **a**. In the process of mitosis, the nuclear membrane undergoes breakdown/reforming cycles and lamin A undergoes depolymerization and polymerization cycles, in which pSer22 plays critical role, resulting in the increase and decrease of FRET ratio in principle. **b**. Time-lapse FRET/CFP ratio images in Hela cells expressing WT LAPS during cell mitosis upon RO-3306 release at t=0. **c**. Time-lapse FRET/CFP ratio images of the WT and R605A in Hela cells as the cell progressed from interphase into mitosis. HeLa cells were released from double thymidine in this experiment. Scale bar = 10 μm. **d**. Time courses of the normalized FRET/CFP ratio of the WT and R605A (n=20, 20) in Hela cells as the cell progressed from interphase into mitosis. **e**. Time-lapse FRET/CFP ratio images of the WT and R605A in Hela cells as the cell progressed from mitosis into interphase. Scale bar=10 μm. **f**. Time courses of the normalized FRET/CFP ratio of the WT and R605A (n=20, 20) in Hela cells as the cell exits mitosis.

To further investigate the pSer22 propagation at nuclear periphery in spatial, we precisely tracked this process with LAPS on confocal microscopy. The results showed that the pSer22 modification on lamin A occurred firstly at the inside of the nuclear membranes, as the phosphorylated portion of lamin A transported into nucleoplasm. With mitosis proceeded, lamin A in the immediate vicinity of the inner nuclear membrane was phosphorylated until the nuclear membrane completely collapses (**Fig 4a**). We profiled the FRET ratio from inside to outer layer of NE, and obviously found that FRET ratio at outer layer of NE was as high as inside when chromatin got condensed into chromosomes, which took about half-an hour to spread out on the NE since cells got released from RO-3306 (**Fig 4a-b**), indicating that pSer phosphorylation on lamin A occurred inside nuclear membrane first to outer layer of NE later.

**Fig 4.**
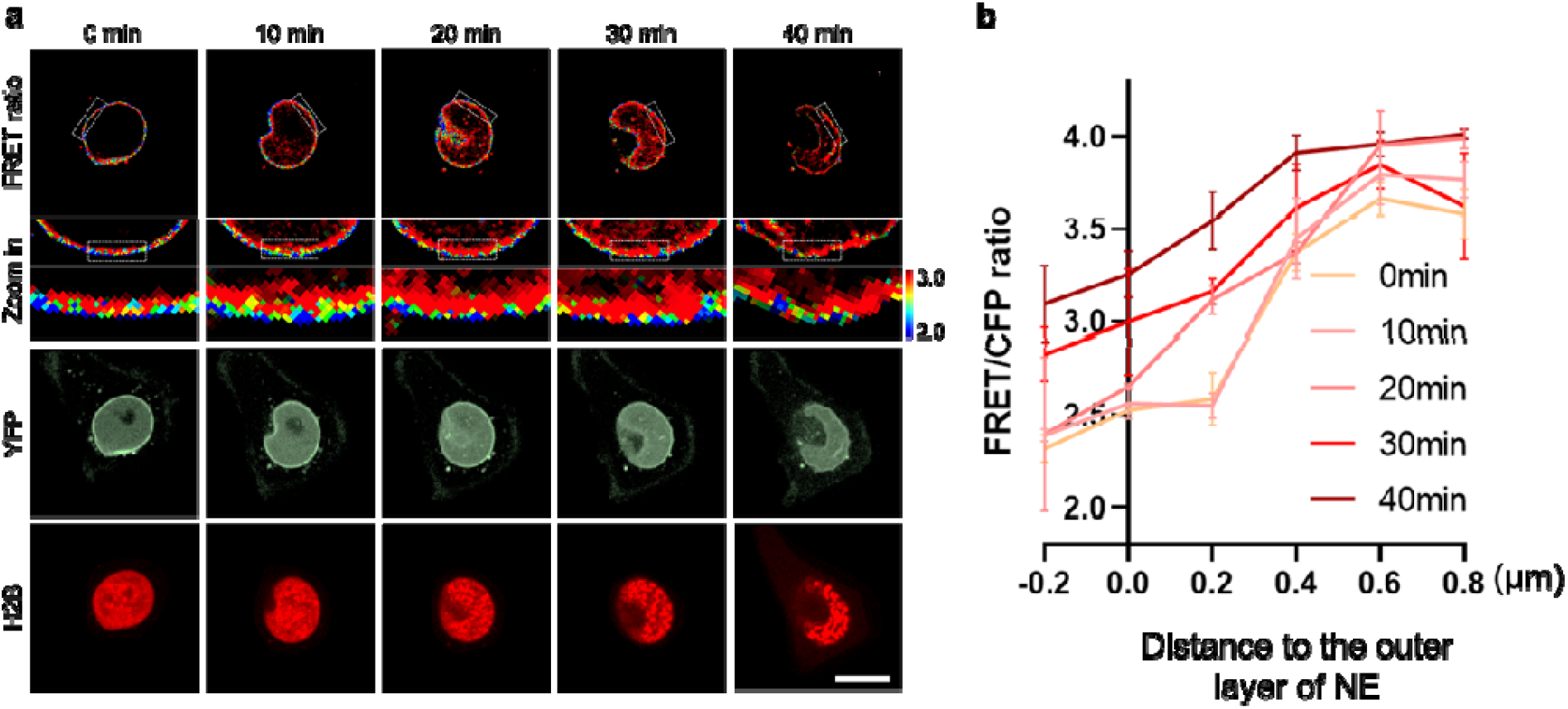
The dynamic and spatial feature of pSer22 at mitosis entry. **a**. Time-lapse imaging was performed in HeLa cells expressing LAPS and H2B-RFP released from RO-3306. FRET ratio: Time-lapse FRET/CFP ratio images, Zoom in: enlarged FRET ratio images. YFP: YFP only channel to show the location of LAPS. H2B: H2B-RFP to indicate chromatin. Scale bar = 10 μm. **b**. Time courses of the FRET/CFP ratio in Hela cells in (a). The abscissa 0 indicates the outer layer of the nuclear membrane. n = 20. The FRET ratio increased nearby 0 μm when cells got into mitosis, indicating the propagation of pSer22.

### A positive correlation between H3S10ph and lamin A pSer22 at mitosis entry

Mitosis is accompanied by a significant increase in both lamin A phosphorylation and H3S10 phosphorylation (H3S10ph), we were curious about which phosphorylation event occurs first and if they have crosstalk each other. To address this, we co-stained pSer22 lamin A and H3S10ph in unsynchronized HeLa cells and did observe pSer22 lamin A and H3S10ph intensity increased dramatically in the mitosis. Interestingly, there was a small portion of H3S10ph in a confined region near the nuclear membrane in some population cells that with moderate level of lamin A phosphorylation (**Fig S3**). To precisely explore the spatiotemporal relationship between the pSer22 lamin A and H3S10ph during cell cycle progression, we synchronized the HeLa cells at G1/S boundary by double thymidine arrest. The immunostaining and WB results demonstrated pSer22 lamin A and H3S10ph gradually increased upon RO-3306 release (**Fig 5a-c**). However, the pSer22 lamin A showed a slight raise in the S phase which might be governed by cell cycle dependent kinase in S phase. Compared with pSer22 lamin A, the pS10 modification on histone 3 was a more specific events in mitosis. Then, we explored the H3S10ph level during mitosis in lamin A knock out cells. The immunostaining and western blot results proved that lamin A loss had no effect on mitosis-induced pSer10 modification on histone 3 (**Fig S4**). To further uncover the crosstalk between pSer22 lamin A and H3S10ph, we inhibited the Aurora B kinase activity by hesperadin^26, 27^ and tracked the pSer22 lamin A dynamics by our LAPS biosensor (**Fig 5d-e**). The results indicated that the mitosis process was delayed and shortened after hesperadin treatment as reported^28^. Meanwhile, the pSer22 on lamin A was also delayed in hesperadin treated group, but the mitosis induced FRET raise was maintained, which implied that Aurora B indeed governed cell mitosis entry, but it was not the key kinase of lamin A. To further explore the delay of mitosis induced by hesperadin, we synchronized the HeLa cells at G2/M boundary by RO-3306 and inhibited the Aurora B activity by hesperadin. The results of western blot indicated that phosphorylation of H3S10 was completely abolished and pSer22 modification on lamin A was prevented, which should be the consequence of delayed mitosis (**Fig 5f-g**), suggesting the positive correlation between H3S10ph and lamin A pSer22 at mitosis entry.

**Fig 5.**
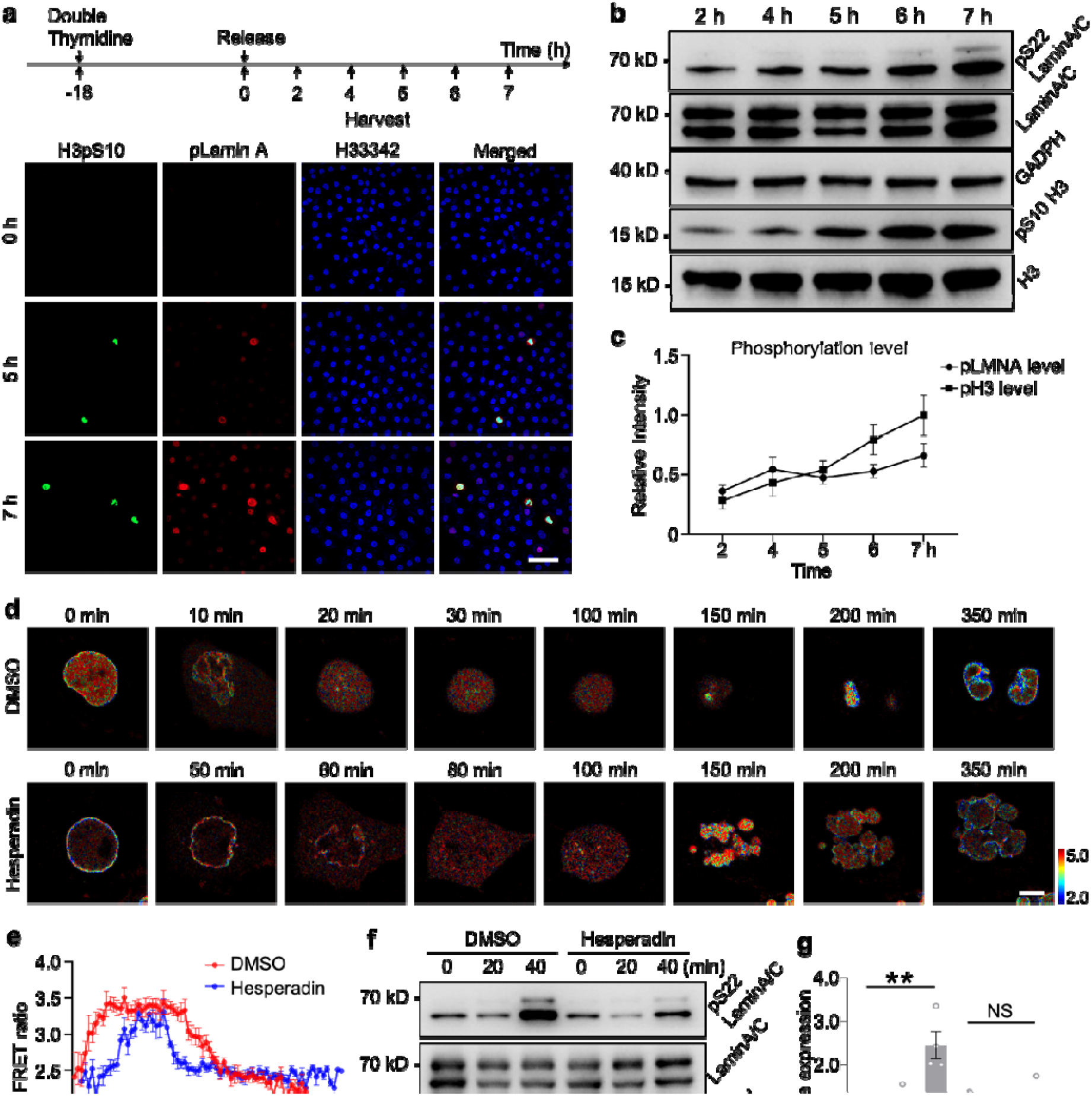
A positive correlation between H3S10ph and lamin A pSer22 at mitosis entry. **a**. Immunofluorescence was used to detect the changes of H3pS10 and pLamin A in HeLa cells from early S phase to M phase after double thymidine release. Green represents H3pS10, red represents pLamin A, and blue represents nucleus. Cells were released after two Thymidine treatments synchronized to the G1/S boundary, and samples were collected at different time points after release, scale bar = 100μm. **b**. WB was used to detect the changes of H3pS10 and pLamin A in HeLa cells from early S phase to M phase, and the cells were synchronized to G1/S boundary after two thymidine treatments and then released at different time points. **c**. WB quantification from (b). **d**. Cells were pretreated with RO-3306 and synchronized to G2/M transition. Time-lapse imaging of WT LAPS in Hela cells treated with DMSO or 200 nM Hesperadin. Scale bar = 10 μm. **e**. Time courses of the normalized FRET/CFP ratio of the Hela cells treated with DMSO or 200 nM Hesperadin in (d) (n=17). **f**. WB was used to detect the changes of H3S10ph and lamin A pSer22 when cells entered mitosis with either DMSO or hesperidin treatment. The cells were synchronized to the G2/M transition after RO-3306 treatment, then collected at 0, 20, and 40 min for WB. **g**. The quantified bar graph for (f) (**P<0.01, not significant (NS)), n = 4.

## Discussion

In summary, we developed a genetically encoded FRET biosensor LAPS for live cell imaging of pSer22 lamin A with high specificity and sensitivity. Utilizing LAPS, we uncovered the gradual reduction of pSer22 modification on lamin A during cell adhesion and hypotonic treatment, which demonstrated how nuclear deformation and cell tension regulates lamin A pSer22. In addition, we monitored the dynamic change of pSer22 lamin A during mitosis with high spatiotemporal resolution. We found that lamin A was phosphorylated from the interior of nuclear membrane to the boundary and was dephosphorylated upon cells exit mitosis. Considering the synchronicity of pSer22 lamin A and H3S10ph, we focused on the crosstalk of these two phosphorylation events. Knockout lamin A didn’t affect H3S10ph in cell mitosis. Inhibition of Aurora B, the kinase of H3S10ph, delayed and shortened the mitotic process while pSer22 modification on lamin A was maintained.

We also observed H3S10ph puncta in very early prometaphase accompanied with moderate pSer22 lamin A, which implied that S10ph modification of histone 3 may start at the nuclear membrane, which eventually prorogate onto entire chromosome. However, so far, we did not work out the simultaneous monitoring of H3S10ph and lamin A pSer22 by dual FRET imaging due to the low sensitivity of H3S10ph biosensor, which was reconstructed with mOrange/Fusion red FRET pair according to previous work^21^. So, a new red FRET pair with high efficiency should be developed in the near future. On the other hand, the FHA2 domain recognizes the phosphorylated threonine in the native circumstance^22^. The affinity of FHA2 domain to phosphorylated serine was relatively low, which impaired the binding efficiency of LAPS. Therefore, novel FHA2 mutants with higher binding affinity to pSer are needed to obtain more dynamic changes of LAPS, which can be optimized by directed evolution of FHA2.

## Acknowledgments

We thank the Bioimaging Core facility at Shenzhen Bay Laboratory for providing imaging support. We also would like to acknowledge engineers Mei Yu and Shixian Huang from the Bioimaging Core facility for assistance with laser-point scanning confocal microscopy (Zeiss 980) and Dragonfly Spinning Disk Confocal Microscopy (7-laser). We thank Yanwei Chen for the help of experimental preparations.

## Author contributions

Q.P., J.Z., and J.H.Q. designed the project and revised the manuscript. J.L. conducted most of the experiments and analyzed data. J.L. and Q.Q.L. wrote the manuscript. J.F.W. analyzed data and organized figures.

## Data Availability

The data that support the findings of this study are available from the corresponding authors upon reasonable request.

## Supplementary Materials

**Fig S1.**
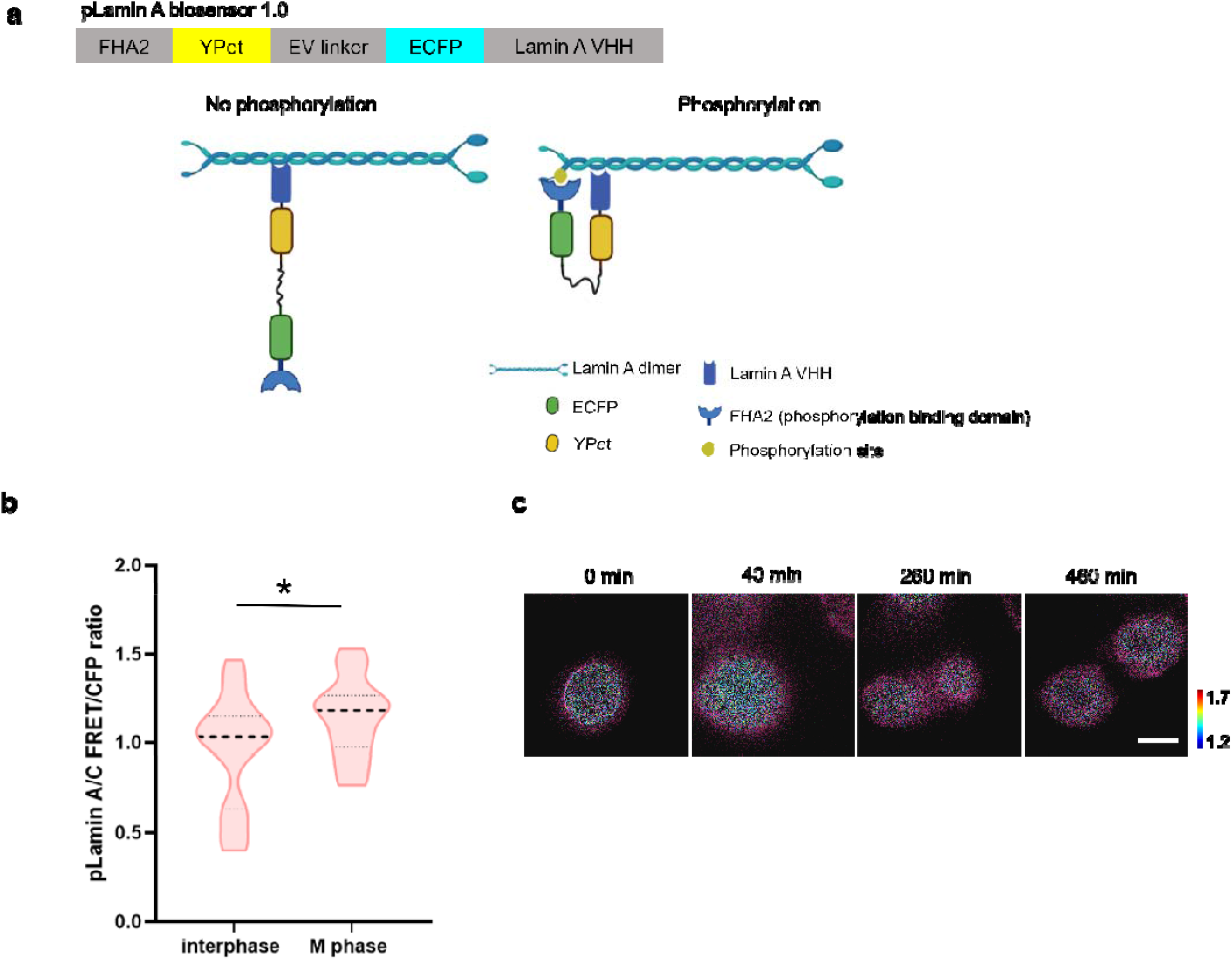
pLaminA biosensor 1.0. **a**. Schematic diagram of the pLaminA FRET biosensor 1.0. A flexible linker (EV linker) with 120 aa was inserted between ECFP and YPet. At resting state when lamin A is not phosphorylated, the excitation of ECFP cannot trigger a FRET event. When lamin A gets phosphorylated, the FHA2 domain can bind to the phosphorylation site, which leads to a tremendous conformational change and proximity of the FRET pair protein. An increase in the FRET ratio could be then observed. **b**. Quantitative results of FRET ratio in HeLa cells expressing pLaminA biosensor 1.0 in interphase and mitotic (M) phase (n = 29, 24). **c**. The time-lapse FRET ratio images of one representative HeLa cells expressing pLaminA biosensor 1.0 in cell mitosis. Scale bar: 10 μm.

**Fig S2.**
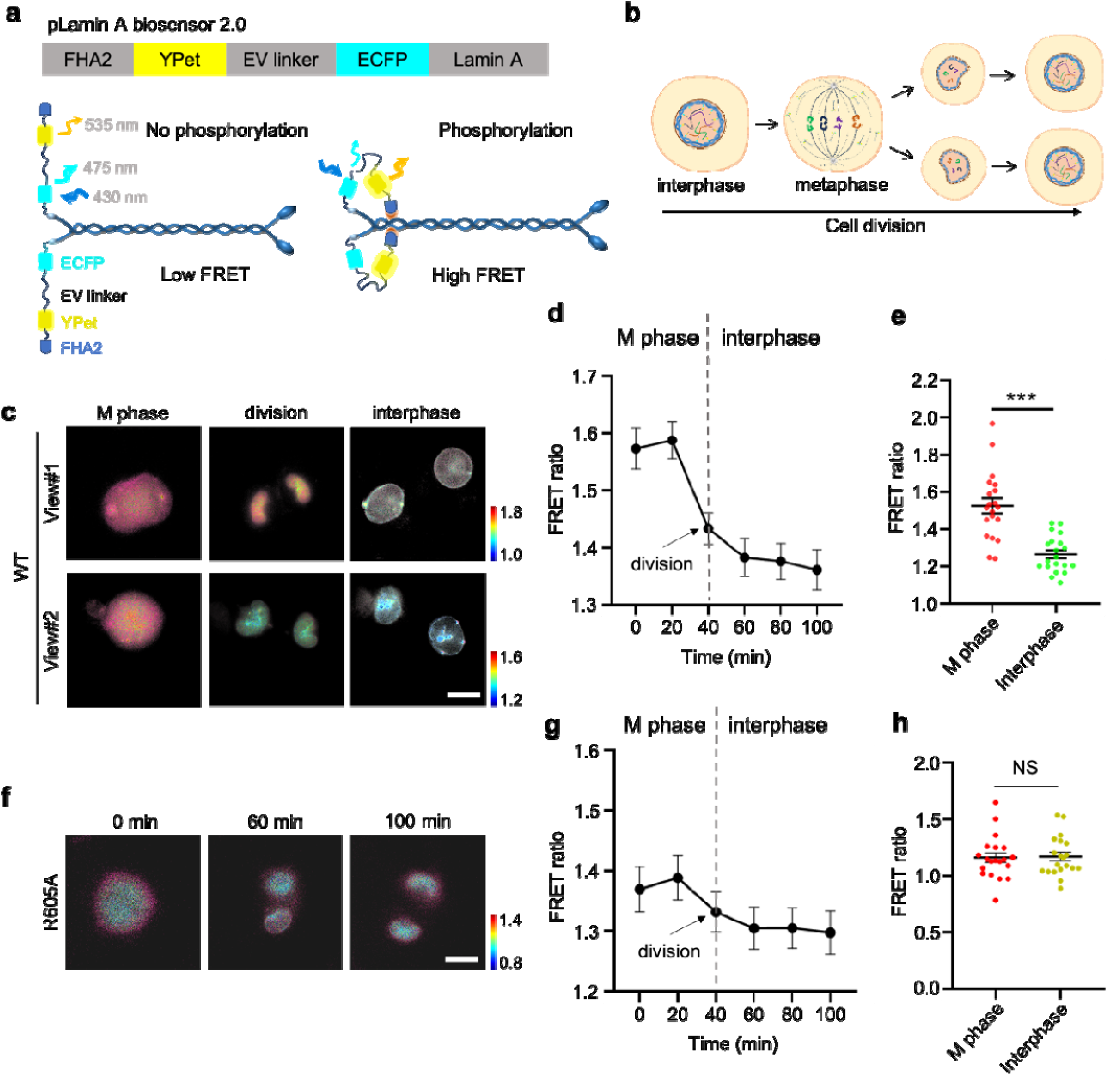
pLaminA biosensor 2.0. **a**. Schematic diagram of the pLaminA FRET biosensor 2.0. A flexible linker (EV linker) with 120 aa was inserted between ECFP and YPet. The Lamin A protein is connected behind the ECFP. At resting state when lamin A is not phosphorylated, the excitation of ECFP cannot trigger a FRET event. When lamin A gets phosphorylated, the FHA2 domain can bind to the phosphorylation site, which leads to a tremendous conformational change and proximity of the FRET pair protein. An increase in the FRET ratio could be then observed. **b**. In the process of mitosis, the nuclear membrane undergoes breakdown/reforming cycles and lamin A undergoes depolymerization and polymerization cycles, in which pSer22 plays critical role, resulting in the increase and decrease of FRET ratio in principle. **c and f**. Time-lapse FRET/CFP ratio images of the pLamin 2.0 biosensor WT and R605A in Hela cells as the cell progressed from mitosis into interphase. Scale bar=10 μm. **d and e**. Quantitative results of FRET ratio in HeLa cells from c. g and h. Quantitative results of FRET ratio in HeLa cells from **f**.

**Fig S3.**
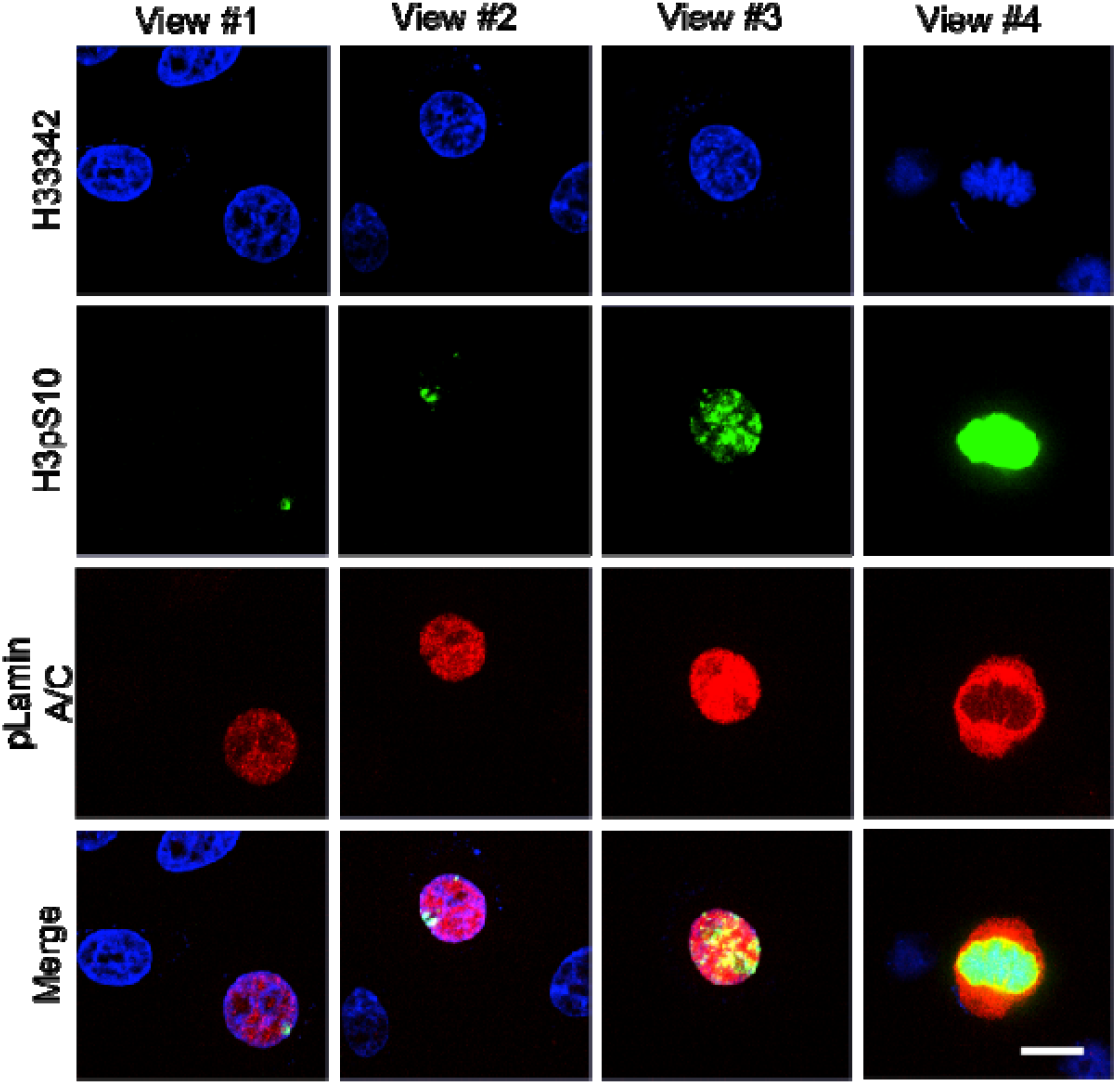
Immunofluorescence images to show H3S10ph and pSer22 Lamin A levels in different view of Hela cells from G2 to M phase. Green represents H3pS10, red represents pLamin A, and blue represents nucleus, scale bar = 10 μm.

**Fig S4.**
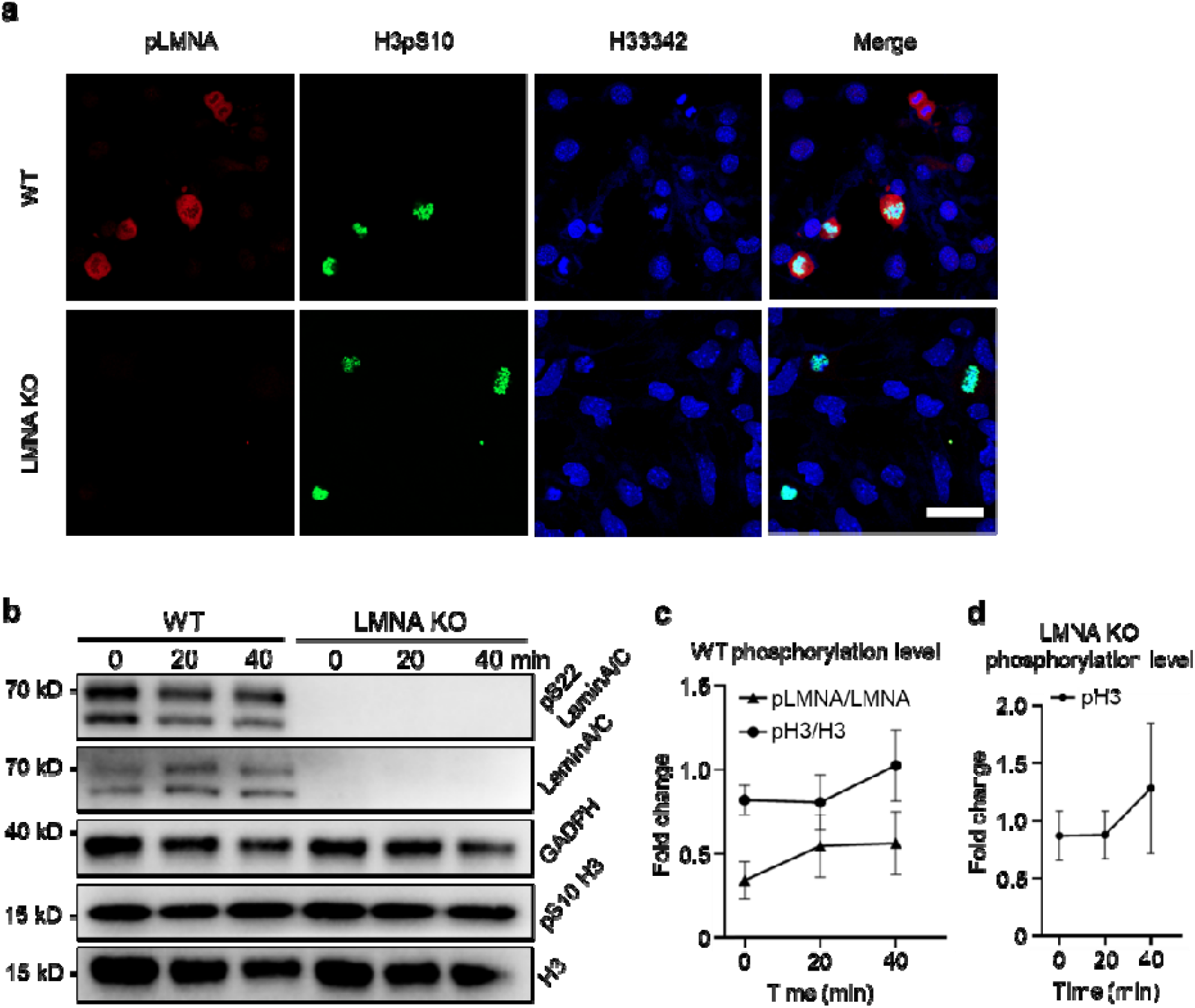
Knockout lamin A had no effect on H3S10ph. **a**. Immunofluorescence was used to detect H3S10ph and pSer 22 Lamin A in WT and LMNA knockout cell lines. Green represents H3S10ph, red represents pSer 22 Lamin A, and blue represents nucleus, scale bar = 50 μm. **b**. WB detected the changes of H3S10ph and pSer 22 Lamin A in WT and LMNA knockout cell lines from G2 to M phase. The cells were synchronized to the G2/M phase after RO-3306 treatment, and the cells were collected at 0, 20, and 40 min for WB detection. **c and d**. WB quantification of both phosphorylation changes in WT cells (c) and LMNA KO cells (d) from b. (n=3)

**Table S1.**
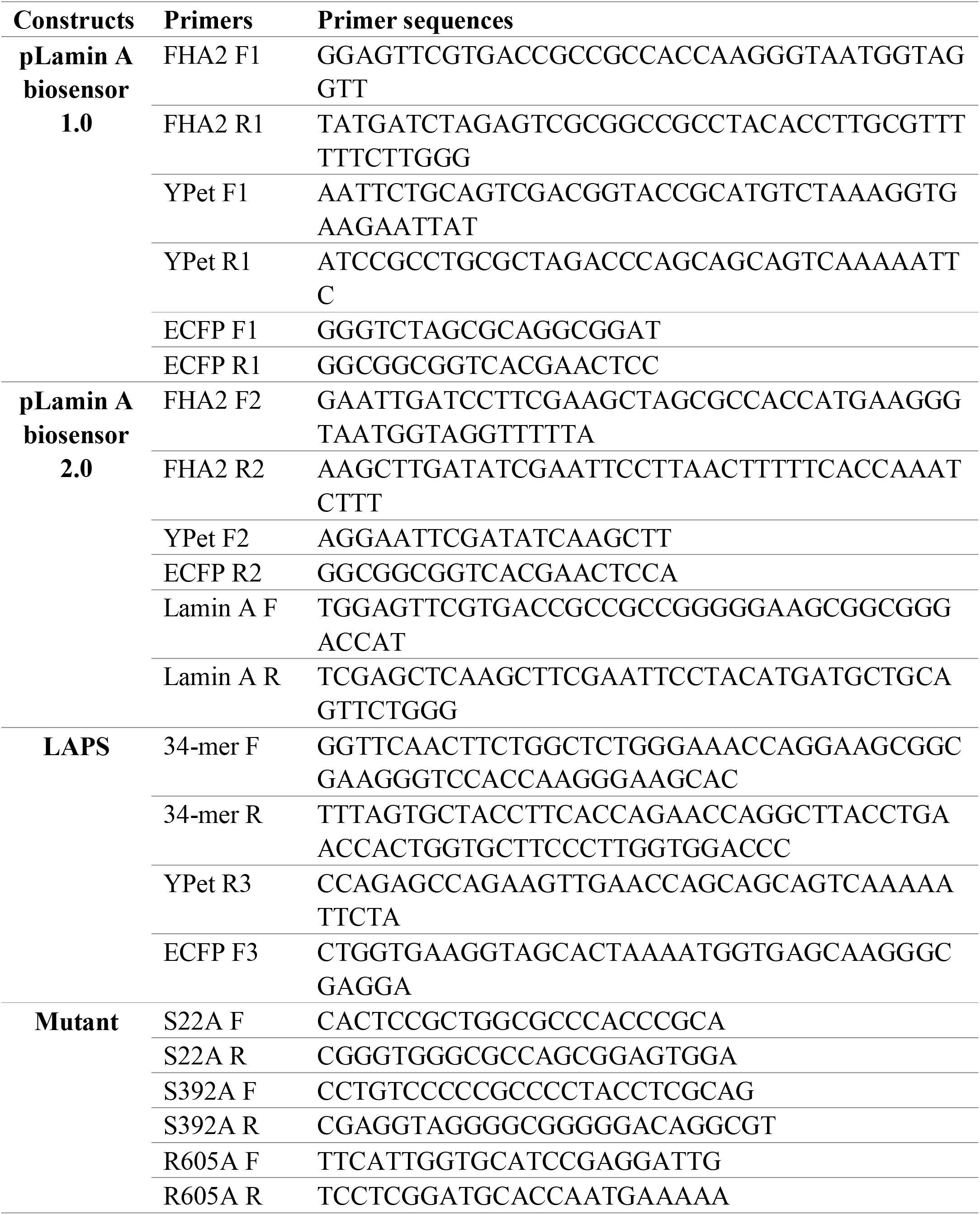
Primers for cloning used in this study.

**Movie S1** (related to Fig 3). Lamin A pSer 22 dynamics during cell mitosis in a representative HeLa cell expressing LAPS.

